# Automatic Recognition of Auditory Brainstem Response Characteristic Waveform based on BiLSTM

**DOI:** 10.1101/2020.10.03.324665

**Authors:** Cheng Chen, Li Zhan, Xiaoxin Pan, Zhiliang Wang, Xiaoyu Guo, Handai Qin, Fen Xiong, Wei Shi, Min Shi, Fei Ji, Qiuju Wang, Ning Yu, Ruoxiu Xiao

**Affiliations:** School of Computer and Communication Engineering, University of Science & Technology Beijing, Beijing 100083, China; College of Otolaryngology Head and Neck Surgery, National Clinical Research Center for Otolaryngologic Diseases, Key Lab of Hearing Science, Ministry of Education, Beijing Key Lab of Hearing Impairment for Prevention and Treatment, Chinese PLA General Hospital, Beijing, China 100853; Institute of Artificial Intelligence, University of Science and Technology Beijing, Beijing 100083, China

**Keywords:** Auditory brainstem response, Characteristic waveform recognition, Neural network model, Bi-directional long short-term memory, Wavelet transform

## Abstract

**Background:** Auditory brainstem response (ABR) test is widely used in newborn hearing screening and hearing disease diagnosis. Identifying and marking are challenging and repetitive tasks because of complex rules of ABR characteristic waveform and interference of background noise.

**Methods:** This study proposes an automatic method to recognize ABR characteristic waveform. First, binarization is created to mark 1024 sampling points accordingly. The selected characteristic area of ABR data is 0-8ms. The marking area is enlarged to expand feature information and reduce marking error. Second, a bi-directional long short-term memory (BiLSTM) network structure is established to improve relevance of sampling points, and an ABR sampling point classifier is obtained by training. Finally, mark points are obtained through thresholding.

**Results:** Specific structure, related parameters, recognition effect, and noise resistance of network were explored in 614 sets of ABR clinical data, and recognition accuracy of waves I, III, and V can reach 92.91%.

**Discussion:** Thus, the proposed method can reduce the repetitive work of doctors and meet accuracy effectively. Therefore, this method has clinical potential.

## 1. Introduction

ABR is an electrical activity of nerve impulses in brainstem auditory conduction pathway caused by acoustic stimulation. It can observe functional status of auditory nerve and lower auditory center, and reflect conduction ability of brainstem auditory pathway [1,2]. Given that patient’s hearing impairment can be diagnosed without his active cooperation, ABR has become one of the routine methods for newborn hearing screening and adult hearing disease diagnosis [3,4,5]. ABR waveform usually has a short eclipse period of 10ms, and electrode intensity in microvolts is recorded. ABR can usually record seven normal phase waves, that are indicated by Roman numerals I, II, III, IV, V, VI, and VII in clinical medicine. Where, waves I, III, and V are often used as clinical diagnosis basis because of their obvious characteristics [6,7]. Figure 1 states the annotated ABR waveforms, which mainly identify waves I, III, and V clinically. Other characteristic waves are usually not displayed clearly because of small amplitude, two-wave fusion and noise interference. Thus, they are rarely used as a basis for diagnosis.

**Fig. 1.**
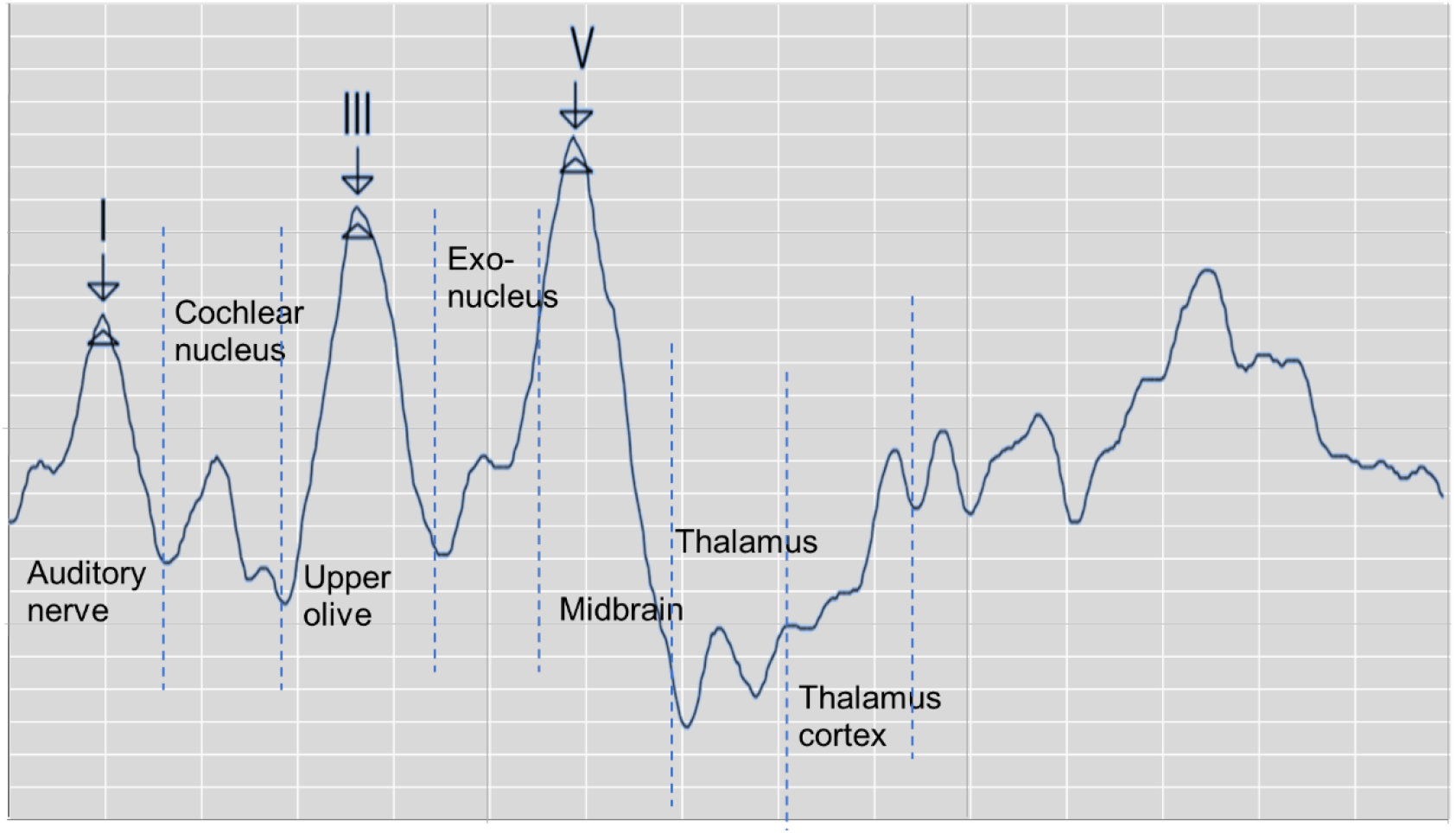
Annotated ABR waveforms

In clinical diagnosis, the minimum short sound stimulation intensity of wave V is usually used as ABR threshold. Sometimes when wave III is greater than wave V, ABR threshold is judged by stimulation intensity of wave III [8]. In determining lesions, the location can be judged according to the eclipse period of waves I, III, and V and the eclipse period between waves and binaural waves [9]. Furthermore, types of deafness of a patient can be judged by observing the change characteristics of ABR waveform latency and special shape of ABR waveform in the same patient under different stimulation levels. Thus, ABR threshold and eclipse period of waves I, III, and V, which is of great significance in clinical applications can be obtained by identifying position of the characteristic wave of ABR. Usually, acquired auditory brainstem evoked potential signal weakened fully. In clinical testing, multiple tests must be performed to superimpose, average, and obtain relatively stable waveform results. This process is susceptible to interference from spontaneous signals. In addition, electrodes placed on the top of skull or mastoid will be disturbed by outside world, thereby resulting in unobvious peaks of evoked potential waveform, overlapping peaks, and false peaks. Performing multiple tests on patients and comparing results, which not only consume a lot of time but are also prone to subjective judgment errors, are usually necessary. Thus, identifying waveform characteristics of ABR and avoiding interference caused by unclear differentiation, fuzzy characteristics, and abnormal waveforms are important issues that needs to be solved urgently and correctly in clinical ABR auscultation.

The application of computer technology in assisting medical diagnosis can effectively reduce errors caused by repetitive work and complex waveform characteristics. This research direction has been important for ABR consultation for a long time [10]. For example, Wilson [11] discussed the relationship between ABR and discrete wavelet transform reconstructed waveforms, indicating that the discrete wavelet transform waveform of ABR can be used as an effective time-frequency representation of normal ABR but with certain limitations. Especially in some cases, the reconstructed ABR discrete wavelet transform wave is missing because of the invariance of discrete wavelet transform shift. Bradly and Wilson [12] further studied the method of using derivative wavelet estimation to automatically analyze ABR, which improved accuracy of main wave identification to a high level. However, they also mentioned the need for further research on the performance of waveform recognition of abnormal subjects, and manual judgment of abnormal waveforms is still required under clinical conditions. Zhang et al. [13] proposed an ABR classification method that combined wavelet transform and Bayesian network to reduce the number of stimulus repetitions and avoid the nerve fatigue of the examinee. Important features are extracted through image thresholding and wavelet transform. Subsequently, features were applied as variables to classify using Bayesian networks. Experimental results show that the ABR data with only 128 repetitive stimulations can achieve an accuracy of 84.17%. Compared with the clinical test that usually requires 2000 repetitions, the detection efficiency of ABR is improved greatly, but the eclipse period is prolonged by 0.1ms. Moreover, the wave intervals of waves I_III and waves III_V tend to be equal. Unfavorable factors will occur when the wave interval is applied as a diagnostic index.

As a heuristic method, neural network can compensate for the lack of feature extraction in traditional methods to improve recognition accuracy, thereby becoming a research direction in recent years [14]. For example, Gholami-Boroujeny et al. [15] proposed a nonlinear adaptive noise cancellation algorithm based on multilayer perceptron neural network, and compared it with a linear adaptive noise cancellation algorithm based on least mean square adaptive filtering. The results show that their method requires less recording time and performs better under low signal-to-noise ratio. Molina et al. [16] proposed a method to classify ABR through symbolic pattern discovery. Initially, the numerical time series is converted into symbolic time series. Then the symbol pattern discovery technology is applied to the output symbol sequence. Finally, a classification technology based on the recognition pattern is used to classify new individuals. The researchers stated that medical personnel accept this method easily because the system will output diagnostic results with medical terminology. Fallatah et al. [17] proposed a new algorithm for detecting speech ABR. The detection time is shortened without reducing the accuracy by using the constructed spectral feature vector as the input of the neural network. In comparison with four methods that are based on wavelet transform and approximate entropy artificial neural network, this method has higher recognition accuracy and only requires a small amount of running time. Although neural networks have been explored in the field of ABR auscultation, they mainly focus on preprocessing or qualitative judgment, and cannot quantify the location of characteristic waveform. Also, the established model cannot fully extract the ABR waveform characteristics. Thus, meeting the accuracy of clinical requirements is difficult. Therefore, artificial neural network is still a very important challenge in identifying ABR waveforms automatically.

In summary, automatic recognition of ABR waveforms through computer-assisted methods can provide diagnostic evidence and assist clinicians and audiologists in ABR interpretation effectively. It also reduces the errors caused by subjective factors, the interference of complex waveforms, and the burden of a large number of repetitive tasks for medical staff. The neural network has long-term research value in the recognition of ABR characteristic waveforms. This study proposes a method of using LSTM network to identify waves I, III, and V in the ABR waveform, and proposes a new idea for the recognition of ABR characteristic waveforms by neural networks. The structure of the study is organized as follows: The experimental data and the detailed description of the proposed method are presented in Chapter 2. Chapters 3 presents the experimental design and the corresponding results. Finally, Chapter 4 discusses this work.

## 2. Materials and Methods

### 2.1 Data Source

The data are provided by Department of Otolaryngology Head and Neck Surgery, Chinese PLA General Hospital. The SmartEP evoked potential test system developed by the American Smart Listening Company is used for measurement and acquisition. Figure 2 shows the clinical collection process where a represents skin degreasing to enhance conductivity; b represents the position of the forehead and earlobe electrodes; c represents the positional relationship diagram of the preamplifier, electrodes, and plug-in earphones; and d shows the details of the preamplifier. The collected waveform is stored in server e and can be observed with the monitor. The clinical short-sound ABR data were collected by 614 subjects at 96dB stimulation intensity after 1024 repeated stimulations. The data contain 1024 sampling points that range from −12.78ms to 12.80ms with an average interval of 0.025ms between every two sampling points. All data were marked by three clinical audiologists with characteristic waves: wave I, wave III, and wave V and cross-validated. Finally, the data were randomly divided into training and test sets. A total of 491 training sets were used to train the network model, and 123 test sets were used for the final recognition accuracy test.

**Fig. 2.**
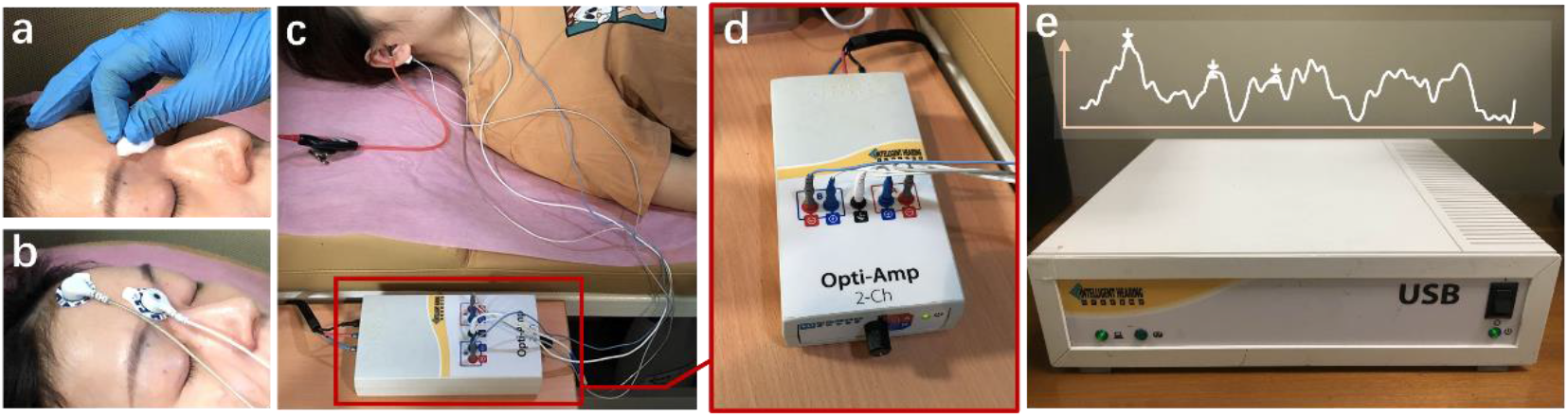
ABR hearing diagnosis clinical collection process. a represents skin degreasing to enhance conductivity; b represents the position of the forehead and earlobe electrodes; c represents the positional relationship diagram of the preamplifier, electrodes, and plug-in earphones; and d shows the details of the preamplifier; The collected waveform is stored in server e and can be observed with the monitor.

### 2.2 Data Processing

To quantify waveform and label points, two 1024×1 matrices were generated as the classification train and label of 1024 sampling points, respectively. The equation of the original training data *A* is expressed as follows:

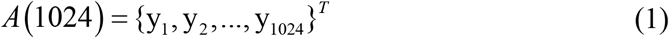

where *y*_*i*_ represents the amplitude of the input ABR data. The position of the serial number corresponds to the position of the ABR data sampling point.

For label *B*, 0 and 1 represent non-feature and feature points, respectively.

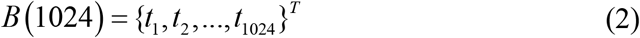

where, *t*_*i*_ is 0 or 1, which indicates whether it is a feature point.

Thus, according to the position of the label value of the label data, the data that correspond to the position of the label matrix was changed to 1 to meet the binary classification requirements of all sampling points. However, abnormal fluctuations are observed in some experimental data (Figure 3). In this ABR clinical test data, the ABR waveform has an unusual increase in the sampling point at the end because of the fluctuation of characteristic waves VI and VII and the result of the external interference. To prevent the interference caused by abnormal data, the data up to 8ms were selected uniformly to identify the characteristic waves.

**Fig. 3.**
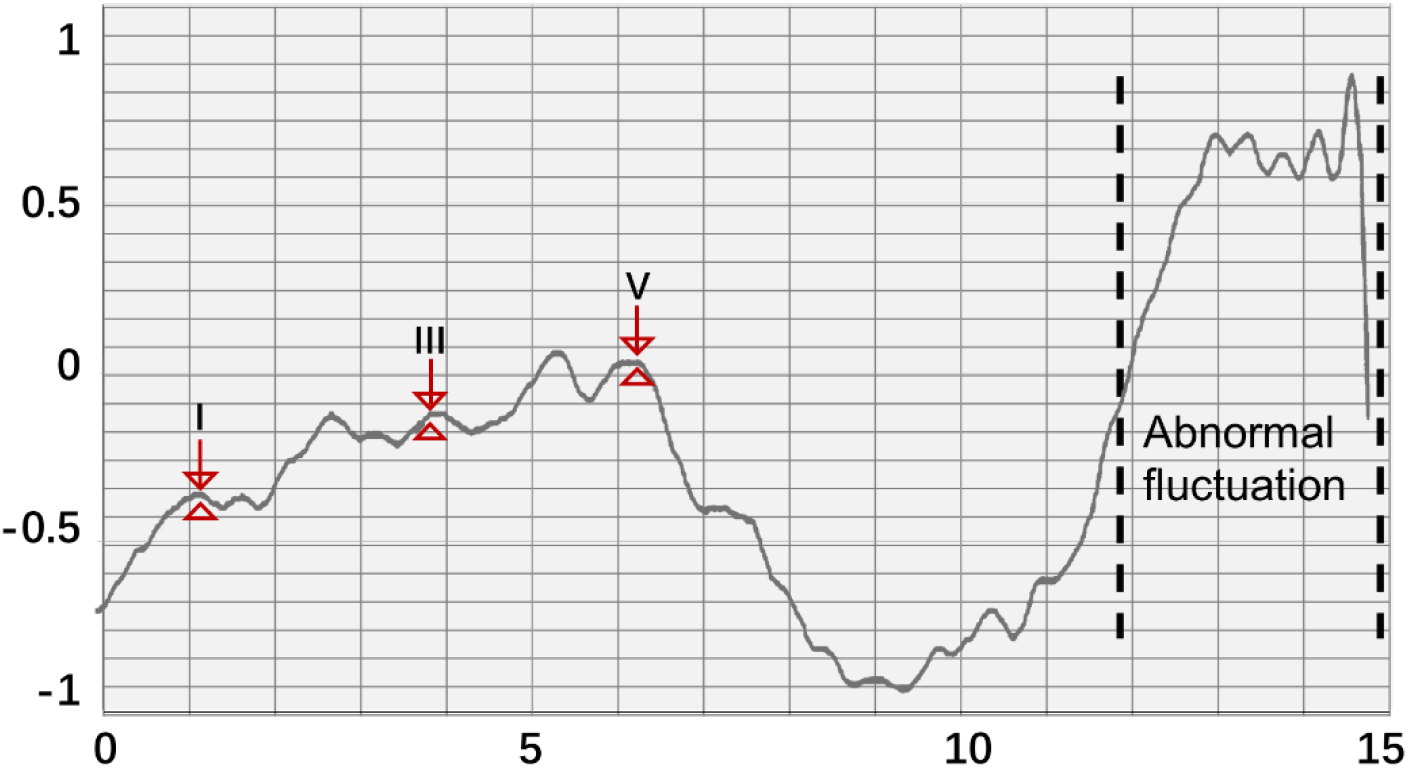
Abnormal ABR waveform and data quantization method

On the other hand, the starting point of the actual stimulation is 0ms. The final potential value input data and the corresponding training label both retained only 321 sampling points of 0-8ms to avoid interference with neural network training and reduce the amount of calculation in the neural network training process. Thus, *A* and *B* are updated as follows:

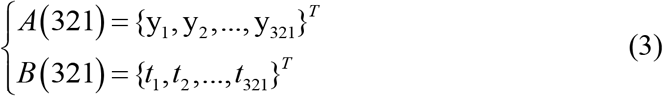

In actual processing, the loss function value can easily reach a low level, and sufficient information cannot be learned because the ratio of the labeled value to the unlabeled value in the 321 sample points is only 3:318. The manually labeled information may also bring certain errors. Thus, this study adopted the method of augmenting the position of the identification point in the training label. The four points (0.1ms) before and after the original marking point were marked as the characteristic area, which expands the marking range of the characteristic waveform.

### 2.3 Network structure

BiLSTM is established as the network structure to enable the input sequence to have a connection with one another [18]. Figure 4 shows that another LSTM layer that propagates backward in time is added on the basis of the unidirectional LSTM forward propagation in time sequence. The final output is determined by the output of the two LSTM layers forward and backward. Compared with the one-way LSTM, the final output avoids the prediction at each time to only be affected by the input of the previous time. Moreover, it can reflect the information characteristics before and after each prediction point better, thereby making more accurate predictions.

**Fig. 4.**
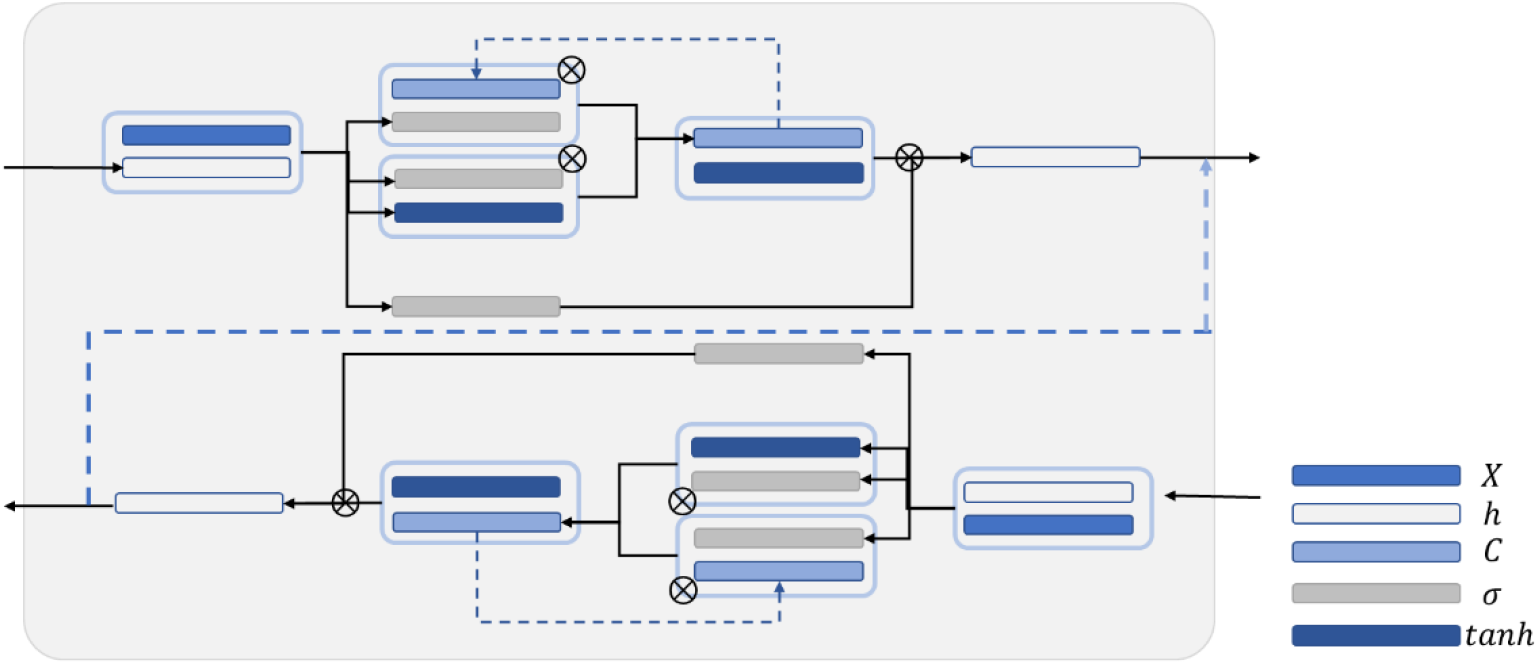
Schematic diagram of BiLSTM structure

The LSTM unit is mainly improved on the basis of the time step unit by adding the output of memory cells to carry information that needs to be transmitted for a long time. Three gate structures are also added. These gate structures are used to select the retention of the memory cell *C*_*t*−1_ value passed from the previous time step, add new information into the memory cell *C*_*t*_, and predict and output the information transmitted by the memory cell and continue to pass it to the next time step.

Figure 5 is a schematic diagram of the LSTM structure. First, to control the proportion of the input information retained by the memory cells at the previous time step, the output *f*_*t*_ is calculated as follows:

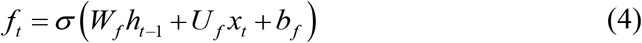

*h*_*t*−1_ is the hidden state value passed at the previous time step; and *W*_*f*_, *U*_*f*_, and *b*_*f*_ are the corresponding weights and biases. The activation function *σ* usually uses the sigmoid function to map the activation value between [0,1]. To control the proportion of information updated into the memory cell, the sigmoid first applied activation function to obtain the output *i*_*t*_. Then, the tanh activation function is applied to obtain *a*_*t*_, and the product of the two is used as the information to update the memory cell. *i*_*t*_ and *a*_*t*_ are calculated as follows:

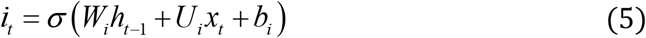

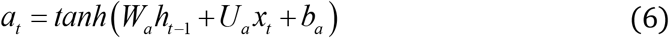

where, *W*_*i*_, *U*_*i*_, *b*_*i*_, *W*_*a*_, *U*_*a*_ and *b*_*a*_ are the weights and biases. Finally, the memory cell *C*_*t*_ is calculated to the next time step by using Eq. (7):

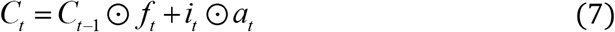

where ⊙ is the Hadamard product, which indicates that the corresponding positions of the matrix are multiplied. The right side refers to the output gate, and the output *o*_*t*_ of the output gate is calculated by using Eq. (8):

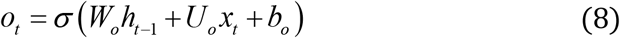

where, *W*_*o*_, *U*_*o*_, and *b*_*o*_ are the weights and offsets. Finally, the output value *h*_*t*_ at the time step is obtained through using Eq. (9):

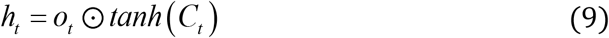

**Fig. 5.**
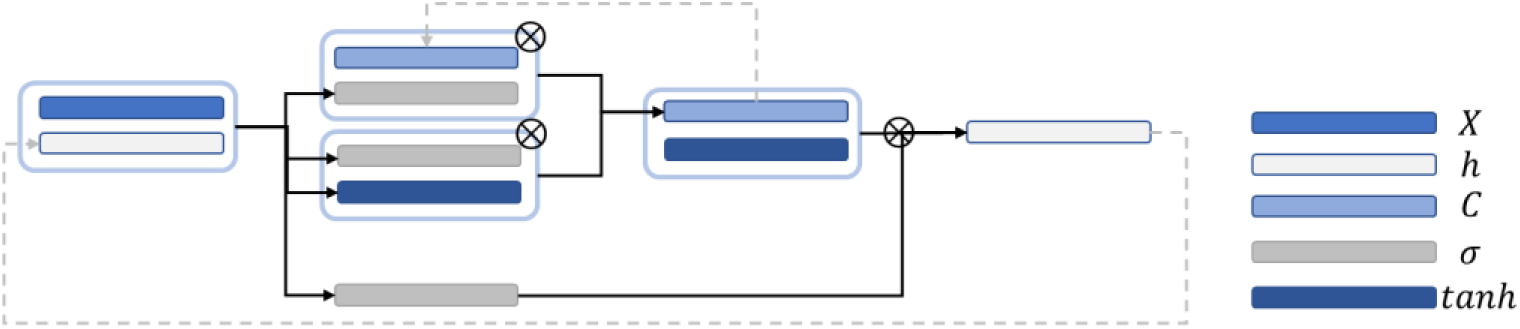
Schematic diagram of LSTM network structure

The predicted output weight *V* and bias *c* are applied to activate the output value to obtain the predicted value 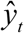, as shown in Eq. (10):

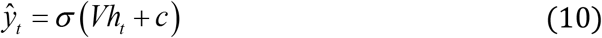

Finally, the loss values 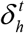 and 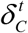 of the hidden state are calculated as follows:

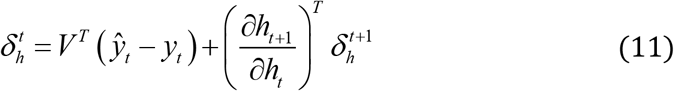

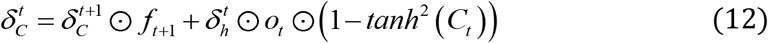

### 2.4 Wavelet Transform

In the traditional mode, wavelet transform is a commonly used method in ABR extraction and recognition research [19]. In ABR extraction, wavelet transform can achieve the effect of eliminating noise by selecting the detail components of specific frequencies for reconstruction and to make the ABR waveform smoother. Obtaining relatively clear waveforms while reducing repetitive stimulation is also possible. Generally, continuous wavelet transform is defined as [20]:

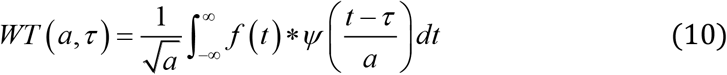

where *f*(*t*) is the signal in the time domain, and the part of 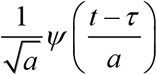 is a wavelet function, which can also be denoted as *ψ*_*a,τ*_(*t*). Two variables, namely, scale *a* and translation *τ*, are available. Scale *a* is applied to control the expansion and contraction of the wavelet function, and the translation amount *τ* controls the translation of the wavelet function. Scale *a* is inversely proportional to its equivalent frequency, which is defined as *φ*(*t*). The complete wavelet expansion:

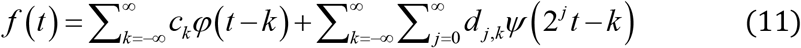

where, *c* and *d* are the coefficients of the corresponding function, *j* is the frequency domain parameter that determines the frequency characteristics of the wavelet, and *k* is the time domain parameter that controls the position of the wavelet base in the time domain. Although the scale and wavelet functions are complex and have different characteristics, the process of wavelet decomposition can be regarded as using a low-pass filter and a high-pass filter to decompose the signal by frequency. The low-frequency components decomposed in each layer are called approximate components, and the high-frequency components are called detailed components.

## 3. Results

In this study, three sets of experiments, including: (1) comparison between various network structures, (2) comparison experiment of wavelet transform, and (3) comparison experiment of different hidden layer nodes were designed. Figure 6 is experimental flowchart. The sequence input layer was used as the input of the potential value of 321 sampling points, and the data were passed to several LSTM or BiLSTM layers. Subsequently, the fully connected layer was connected. The classification probability of each time point was calculated using the softmax function. Finally, the classification layer was connected. The cross-entropy function [21] was used to calculate the loss function of each time point and the overall loss function of the sequence. Then, time sequence was classified.

**Fig. 6.**
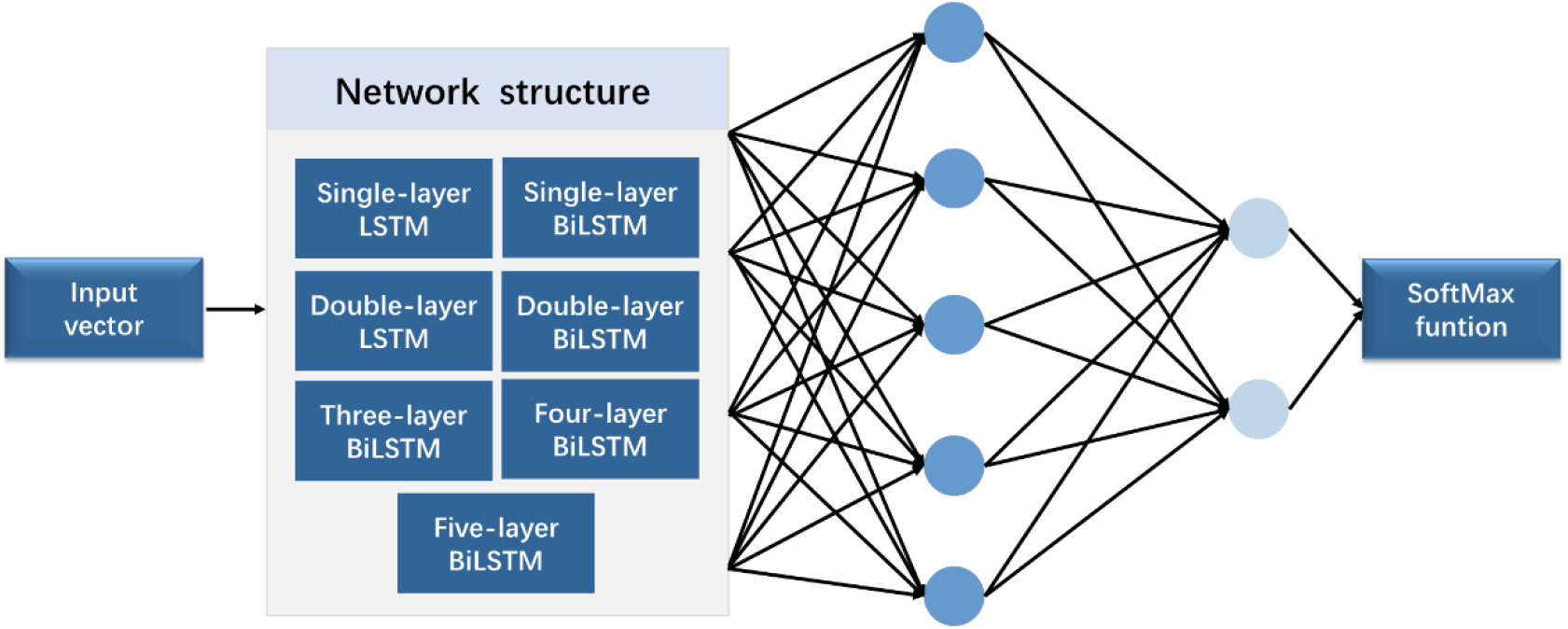
Experimental flowchart.

In the comparison experiment of multiple network structures, seven network structures, including single-layer LSTM, (2) double-layer LSTM, (3) single-layer BiLSTM, (4) double-layer BiLSTM, (5) three-layer BiLSTM, (6) four-layer BiLSTM, and (7) five-layer BiLSTM network layers were selected. In the comparative experiment of different hidden layer nodes, a three-layer bidirectional LSTM network was used for training, and different numbers of hidden neurons were applied. The experiment applied four groups of different numbers of hidden neurons, including: 64, 128, 256, and 512.

In the comparative experiment of the wavelet transform, all data added noise as interference. Seven different network structures were used for testing. For instance, the training data preprocessed by wavelet transform were used as the experimental group, and the training data trained using the original data were used as the control group. In this experiment, ABR data were decomposed in six layers, and the approximate and detailed components of the 6th and 4th, 5th, and 6th layers were retained to reconstruct the waveform, respectively. The parameter configuration is consistent. The network was trained with 5 K-fold cross-validation (K=9), and the test was performed to obtain the average value.

The output results are in the form of ‘region’. Figure 7 expresses the output visualization. Where the curve is the original ABR used for identification, and the red labels are the network prediction classification results reduced by four times. ABR of the first 8ms is clearly divided into two different areas. The part with 0 is the identified non-characteristic wave, and the other part is the identified characteristic wave. A total of 20 sampling points (0.5ms) are set as the threshold: the area within 20 sampling points between the beginning and the end is the same characteristic wave area. Finally, the time mean value of the first and last points is calculated as the time value of the recognized characteristic wave. The similar sampling points are calculated to obtain the unique characteristic wave value. Finally, the recognition accuracy rate is calculated according to the identified ABR feature wave position.

**Fig. 7.**
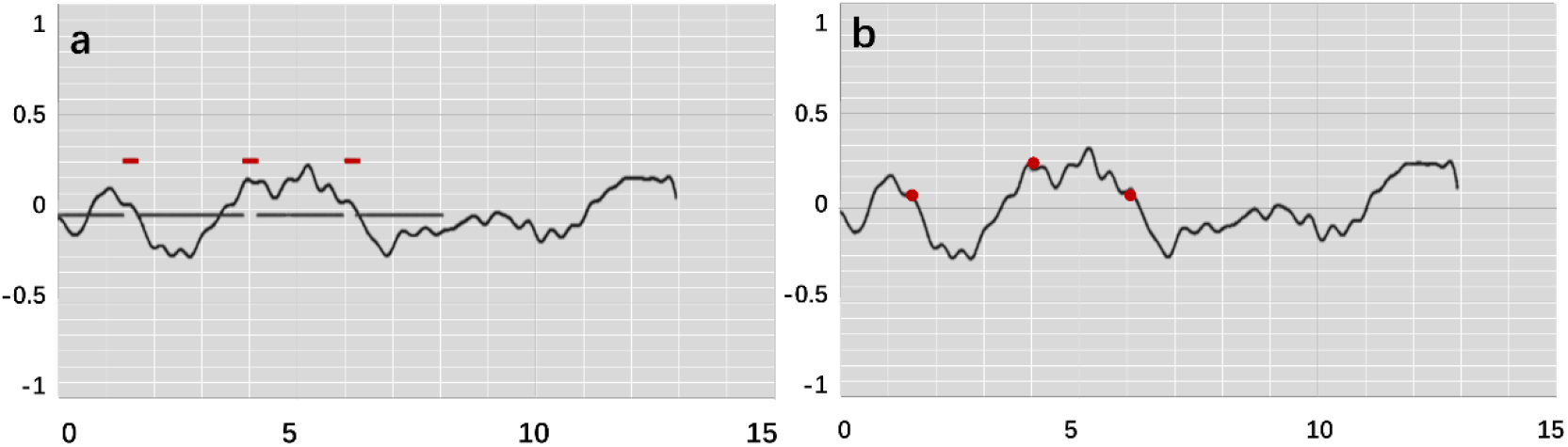
Feature labeling on the ABR

Four recognition results of ABR data were randomly selected and presented in Figure 8. After threshold processing, output vectors from models were converted to feature points. Compared with manual labels, feature points are similar with them in position. These recognitions have clinical potential. Therefore, they also verify possibility of the proposed method. To better verify accuracy of recognition, this work has carried out a quantitative discussion from different network structures, wavelet transform processing and number of hidden neurons.

**Fig. 8.**
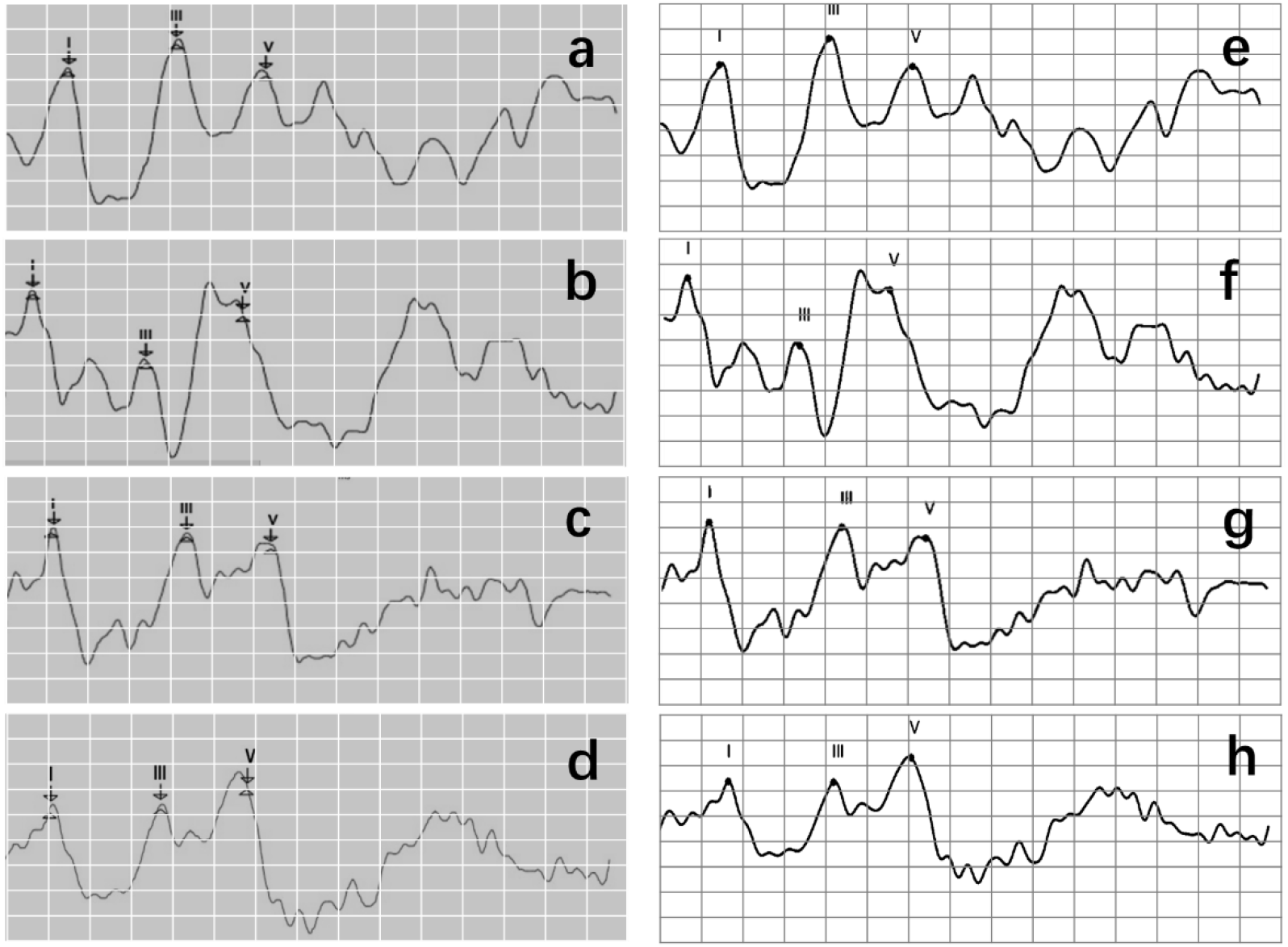
Recognition results of four data, where a, b, c, and d are manual labels. Also, e, f, g, and h represent outputs of model.

### 3.1 Comparison between multiple network structures

Generally, an error scale of 0.2ms is applied as scale range of clinically marked points. The maximum allowable error value ME was set. By traversing all the identification points, if the distance between the eclipse period of the prediction point and the identification point was within the range of ME, then the prediction result was correct. According to the number of correct prediction points *r*_*p*_ and the total marked points *p*_*n*_, the accuracy rate ACC is calculated using *r*_*p*_ / *p*_*n*_, as shown in Eq. (9):

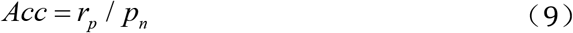

In this study, three error scales ME of 0.1, 0.15, and 0.2ms were calculated respectively to further explore the recognition accuracy and other related laws. Loss value of training results with different network structures and ACC under different error scales are revealed in Table 1:

**Table 1.** Loss value and ACC of each network structure

| Network Structure | Training Loss | Validation Loss | Accuracy (0.1ms) | Accuracy (0.15ms) | Accuracy (0.2ms) |
| --- | --- | --- | --- | --- | --- |
| LSTM | 0.1463 | 0.1635 | 37.08% | 44.92% | 50.37% |
| LSTMx2 | 0.1123 | 0.1625 | 58.61% | 65.75% | 70.59% |
| BiLSTM | 0.1264 | 0.1562 | 61.96% | 72.03% | 77.60% |
| BiLSTMx2 | 0.0849 | 0.1285 | 78.74% | 84.88% | 86.84% |
| **BiLSTMx3** | **0.0704** | **0.1275** | **85.46%** | **91.06%** | **92.91%** |
| BiLSTMx4 | 0.0651 | 0.1342 | 82.48% | 88.32% | 90.20% |
| BiLSTMx5 | 0.0617 | 0.1467 | 83.31% | 88.80% | 90.90% |

Figure 9a indicates data distribution to observe correlation with different network structures visually. Notably, ACC of BiLSTM network is higher than that of LSTM network. In addition, ACC of single-layer BiLSTM network and double-layer LSTM network is similar. The reason is because two-way LSTM network has a similar structure to double-layer LSTM network. However, information in BiLSTM network has characteristics of propagating in positive and reverse directions, whereas two-layer LSTM network only propagates in the positive sequence over time. This phenomenon leads to differences in ACC between the two models. LSTM and BiLSTM networks increase ACC with number of superimposed layers. After BiLSTM network reaches three layers, ACC will no longer increase significantly. Network structure will gradually reach an over-fitting state and increase computational pressure because of excessive parameters. Thus, three-layer BiLSTM network is a better choice.

**Fig. 9.**
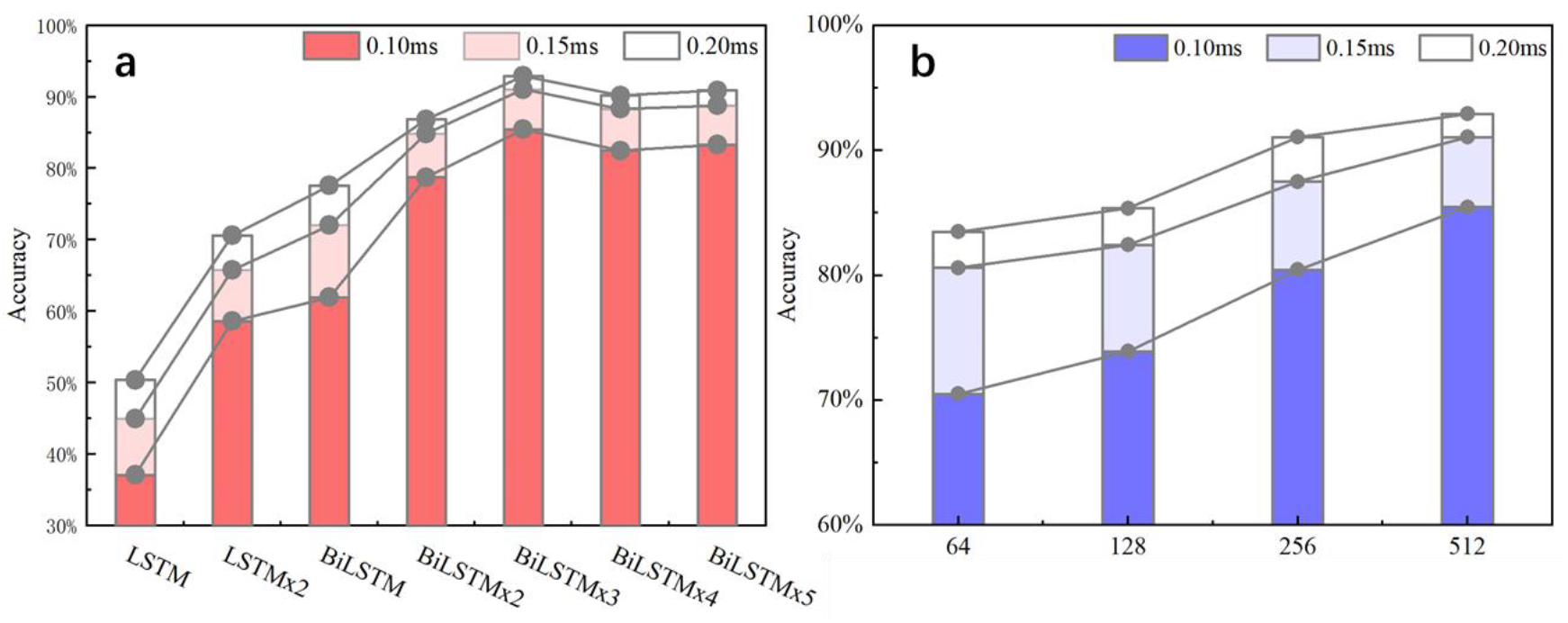
ACC of different network structures and different hidden nodes

### 3.2 Wavelet transform experiment

When testing ACC of wavelet transform, ABR data was decomposed in 6 layers. Also, approximate components of 6th layer and detailed components of the 4th, 5th, and 6th layers were retained to reconstruct waveform. Figure 10 expressed an instance of filtered result by wavelet transform. Curve processed by wavelet transform becomes smoother. Then, unprocessed ABR data served as a control experiment. In this work, detection and comparison were carried out based on two error scales of 0.1 and 0.2ms (Table 2). Results of recognition ACC is expressed in Figure 11:

**Table 2.** ACC of each network structure with original data and wavelet transform data

| Network Structure | Original data (0.1ms) | Wavelet transform data (0.1ms) | Original data (0.2ms) | Wavelet transform data (0.2ms) |
| --- | --- | --- | --- | --- |
| LSTM | **37.08%** | 37.95% | **50.37%** | 52.94% |
| LSTMx2 | **58.61%** | 55.47% | **70.59%** | 72.46% |
| BiLSTM | **61.96%** | 59.17% | **77.60%** | 76.25% |
| BiLSTMx2 | **78.74%** | 73.03% | **86.84%** | 84.71% |
| BiLSTMx3 | **85.46%** | 79.00% | **92.91%** | 90.50% |
| BiLSTMx4 | **82.48%** | 77.73% | **90.20%** | 89.67% |
| BiLSTMx5 | **83.31%** | 78.09% | **90.90%** | 89.17% |

**Fig. 10.**
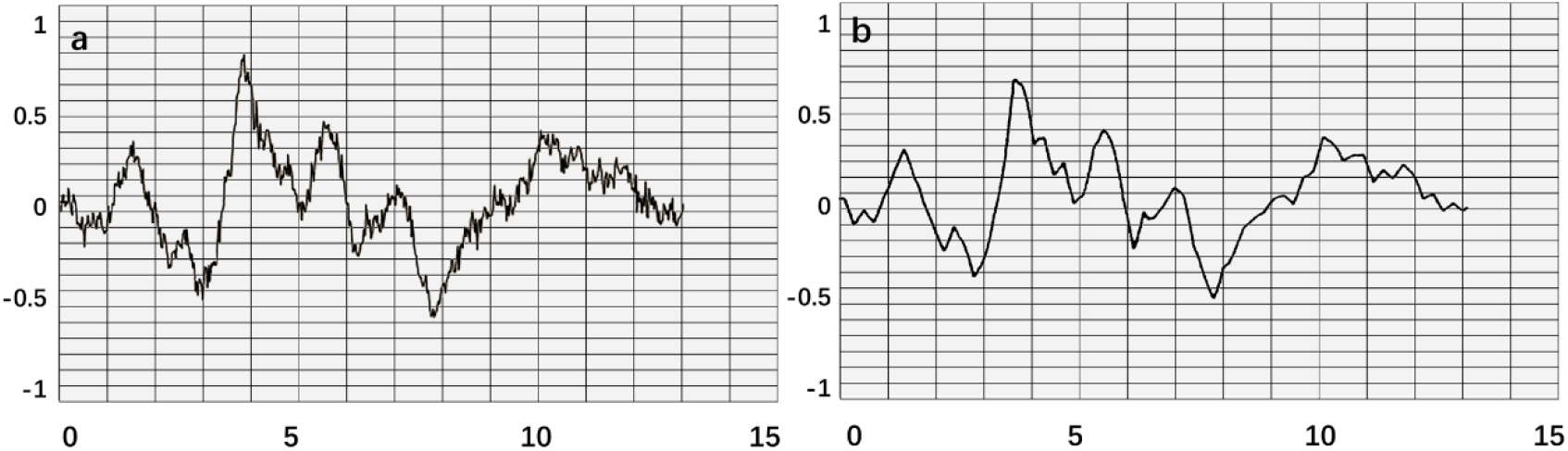
An instance result from wavelet transform, where a is original data. Obvious interference occurred in this waveform. b is obtained after smoothing.

**Fig. 11.**
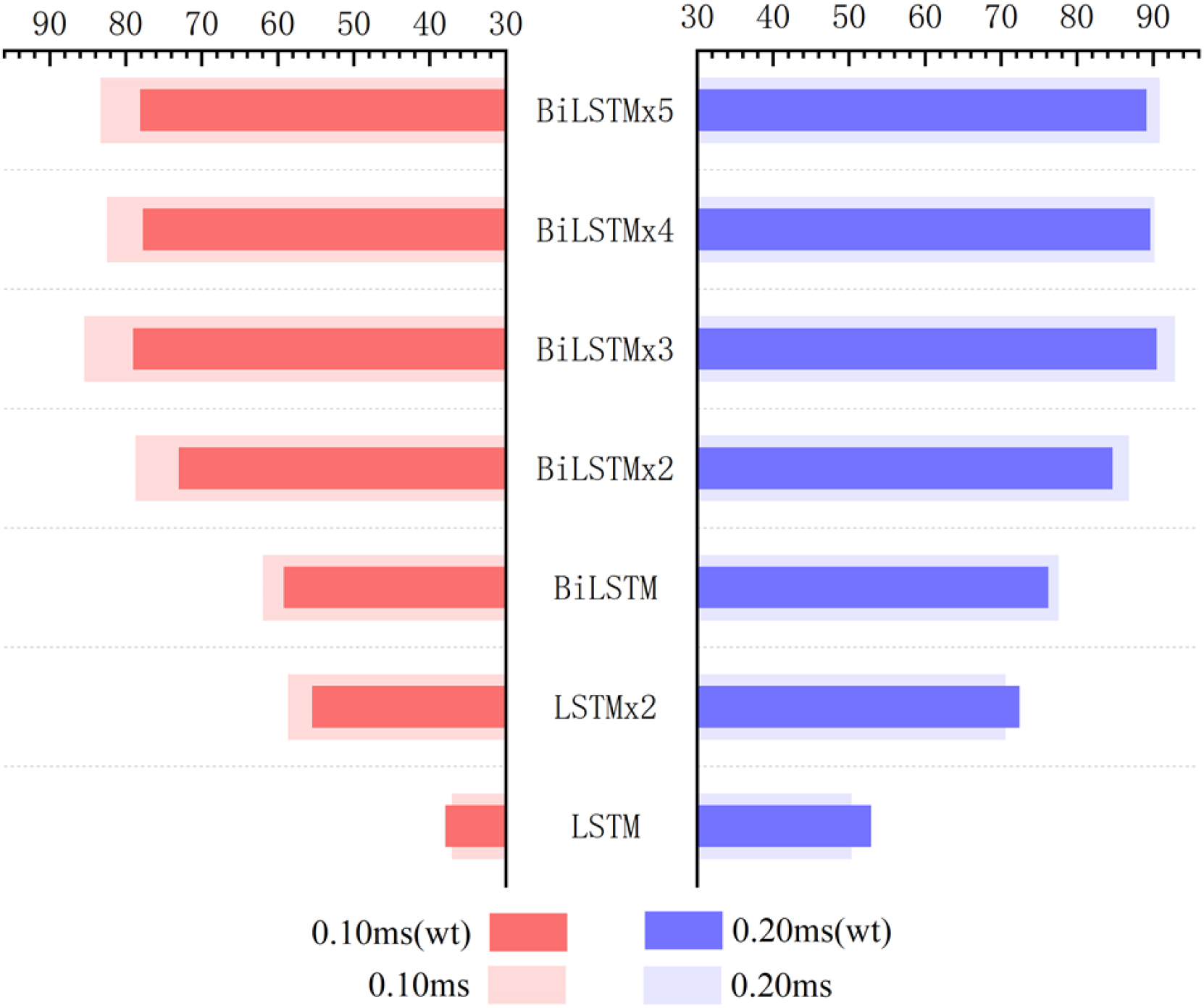
Influence of wavelet transform preprocessing on accuracy. wt represents the results obtained by wavelet transform preprocessing.

Recognition ACC of preprocessing in LSTM network using wavelet transform are slightly higher than that of control group. However, they are not as good as control group in the BiLSTM network. Especially, the highest ACC difference reaches to 6.46% when calculated with a 0.1ms error scale. Also, the difference reduces to less than 3% when calculated with a 0.2ms error scale. Results indicate that wavelet transform preprocessing does not obtain a higher ACC by smoothing curves. Due to wavelet decomposition and reconstruction, a slight deviation was created in the position of wave crest. Some information was destroyed in the ABR waveform, therefore results of training and recognition were affected. This means that BiLSTM network has noise immunity and can handle low-quality ABR data.

### 3.3 Comparative experiments of different hidden layer nodes

Based on above results, three-layer BiLSTM network is a better choice. ACC with different hidden node numbers were discussed in this work (Table 3). Figure 9b expressed ACC results with different hidden layer nodes of 64,128,256 and 512. Obviously, recognition ACC increases with number of hidden nodes, because enough parameters make network fitting accurately. Also, ACC of 0.2ms error scale increases slowly during the change process of 256-512 nodes and has basically saturated. Considering accuracy standard in practical applications and time cost of training that may be brought by increasing number of hidden nodes, a network of 512 hidden nodes is a better choice.

**Table 3.** ACC with different hidden layer nodes

| Hidden layer Nodes | Accuracy (0.1ms) | Accuracy (0.15ms) | Accuracy (0.2ms) |
| --- | --- | --- | --- |
| 64 | 70.50% | 80.61% | 83.48% |
| 128 | 73.90% | 82.44% | 85.36% |
| 126 | 80.44% | 87.49% | 91.07% |
| 512 | 85.46% | 91.06% | 92.91% |

Furthermore, this work mainly discusses characteristic wave recognition process of a short-sound ABR with a 96dB stimulus. Also, only parameters such as latency and wave interval can be obtained. In clinical applications, many indicators can still be used as a diagnostic basis, such as relationship between potential values of different stimulus sizes, response and disappearance of wave V and change of eclipse period of each characteristic wave. This also provide a new idea for the subsequent computer-assisted ABR diagnosis and treatment.

## 4. Discussion

This work proposes an automatic recognition method for ABR characteristic waveforms using BiLSTM network. The main purpose is to identify positions of characteristic waves I, III, and V, which assist medical staff in obtaining relevant clinical test parameters, such as eclipse period and wave interval. A data quantification process is designed to analyze the characteristic waveform of ABR, including selection area of potential signal and expansion of label position. Optimal network model structure is obtained through multiple sets of comparative experiments. In 614 sets of clinically collected ABR waveform experiments, network’s overall recognition of characteristic waves showed an ACC of 92.91%.

Experimental results express that the method proposes a new idea for identification of ABR characteristic waveforms, and helps professionals to obtain eclipse period parameters in ABR waveforms. Therefore, computer automatic identification method can obtain deeper information, avoid subjective judgment error of medical staff in the manual identification process effectively, reduce number of repeated stimulations during test and also avoid the vision fatigue of the tested person. Because of noise immunity of proposed network model, it can effectively reduce repetitive detection of patients. In process of large-scale identification, average time of each data by using the method only takes approximately 0.05s, which is much faster than speed of manual identification. Thus, it has great advantages in repeatable work.

## Author Contributions

Conceptualization, Cheng Chen and Li Zhan.; methodology, Cheng Chen; software, Xiaoxin Pan; validation, Handai Qin, Fen Xiong and Wei Shi; formal analysis, Min Shi; investigation, Fei Ji.; resources, Qiuju Wang; data curation, Xiaoxin Pan; writing—original draft preparation, Cheng Chen and Li Zhan; writing— review and editing, Ruoxiu Xiao and Ning yu; visualization, Li Zhan; supervision, Ning Yu.; project administration, Zhiliang Wang and Xiaoyu Guo; funding acquisition, Ruoxiu Xiao. All authors have read and agreed to the published version of the manuscript.

## Funding

This work was funded by the National Natural Science Foundation of China (61701022), National Key Research and Development Program (2017YFB1002804, 2016YFC0901304), PLA General Hospital (QNC19051), the Active Health Project of the Ministry of Science and Technology (2020YFC2004001), the Fundamental Research Funds for the Central Universities (FRF-BD-20-11A), and the Beijing Top Discipline for Artificial Intelligent Science and Engineering, University of Science and Technology Beijing.

## Conflicts of Interest

The authors declare no conflict of interest. The funder had no role in the design, execution, interpretation, or writing of the study.

## Notes

### Competing Interest Statement

The authors have declared no competing interest.

